# Differential Treatment Benefit Prediction For Treatment Selection in Depression: A Deep Learning Analysis of STAR*D and CO-MED Data

**DOI:** 10.1101/679779

**Authors:** Joseph Mehltretter, Robert Fratila, David Benrimoh, Adam Kapelner, Kelly Perlman, Emily Snook, Sonia Israel, Marc Miresco, Gustavo Turecki

## Abstract

**Background:** Depression affects one in nine people, but treatment response rates remain low. There is significant potential in the use of computational modelling techniques to predict individual patient responses and thus provide more personalized treatment. Deep learning is a promising computational technique that can be used for differential treatment selection based on predicted remission probability.

**Methods:** Using STAR*D and CO-MED trial data, we employed deep neural networks to predict remission after feature selection. Differential treatment benefit was estimated in terms of improvement of population remission rates after application of the model for treatment selection using both naive and conservative approaches. The naïve approach assessed population remission rate in five sets of 200 patients held apart from the training set; the conservative approach used bootstrapping for sample generation and focused on population remission rate for patients who actually received the drug predicted by the model compared to the general population.

**Results:** Our deep learning model predicted remission in a pooled CO-MED/STAR*D dataset (including four treatments) with an AUC of 0.69 using 17 input features. Our naive analysis showed an improvement of remission of over 30% (from a 34.33% population remission rate to 46.12%). Our conservative analysis showed a 7.2% improvement in population remission rate (p= 0.01, C.I. 2.48% ± .5%).

**Conclusion:** Our model serves as proof-of-concept that deep learning has utility in differential prediction of antidepressant response when selecting from a number of treatment options. These models may have significant real-world clinical implications.

## 1 Introduction

Major Depressive Disorder (MDD) is the greatest cause of disability-adjusted life-years (DALYs) lost globally and affects over 300 million people at any given time (World Health Organization, 2017). MDD strongly associates with suicide (Turecki and Brent, 2016), early mortality (Saint-Onge et al., 2014), and represents a significant cost to patients, families, healthcare systems and the economy (Kessler 2012; Stewart et al, 2003). While clinical guidelines (Kennedy et al., 2016) define the optimal treatment outcome for depression to be full remission of symptoms, many patients will not reach remission after a first or second antidepressant treatment (Rush et al., 2006).

There are many different interventions employed by professionals to treat depression, but little is known about which patients will respond best to which treatments, and many possible predictors exist (Perlman et. al, 2019). In this work, we explored the promise of using machine learning techniques to develop personalized medicine approaches to MDD treatment to improve remission rates. We will elaborate on an approach that allows for not only prediction of remission, but the selection of an optimal treatment from a set of options; we call this *differential prediction*.

Machine learning work in this domain has previously borne fruit (see Lin et al., 2018; Iniesta et al., 2016, Lee et al., 2019). Chekroud et al. (2016) used machine learning to predict remission for patients administered citalopram in the STAR*D study with roughly 65% accuracy using only clinical and demographic variables. Iniesta et al. (2018) used machine learning to predict remission in the GENDEP study using clinical, demographic, and genetic information.

In addition to predicting remission, there would be great clinical utility in models that help us assign patients to treatments in a way that improves their chances to remit. Work has also been done in this area, with Derubeis et al. (2014) producing the Personalized Advantage Index (PAI) which can help determine if patients are more likely to benefit from one treatment over another. In addition, specific alleles of genetic polymorphisms mediating processes such as stress response, immune regulation, and neurotransmission were relevant for predicting response to different antidepressants (Uher, 2011).

Most work to date has focused on predicting whether a patient would benefit more from one of two treatment options. However, personalized medicine models must be able to assess *differential* benefit between *multiple* treatments. Should they be able to do so using maximally accessible information, such as simple clinical and demographic information, this would be a significant benefit for populations who cannot yet access genetic testing and other biomarker collection technologies. In addition, modeling procedures should be flexible enough to accommodate additional multimodal data to take advantage of potential biomarkers. Finally, useful personalized medicine models must produce interpretable results - clinicians must be able to understand which factors are significant for the algorithm when it generates predictions.

Deep learning is a powerful machine learning technique based on artificial neural networks that can use multimodal data. This technique has recently gained popularity due to its superior predictive performance on various classification tasks such as image recognition (Goodfellow et al., 2016). We set out to use deep learning to analyze data from two well-known datasets, Sequenced Treatment Alternatives to Relieve Depression (STAR*D) and Combining Medications to Enhance Depression Outcomes (CO-MED), to produce a novel differential treatment selection model that could help select between more than two treatments. Crucially, we used recently developed techniques to validate this model on data unseen by the algorithm to get a sense of what clinical benefit in terms of population remission rates could be expected from its use and found that an algorithm using simple clinical and demographic data could have a significant impact on population remission rates. We chose to create one model in order to accomplish differential prediction, instead of having a separate model for each drug. This was to ensure that we were truly capturing a differential prediction, and not a prediction of general treatment response. We also chose not to create models specific to each drug as we wished to be able to capture differences between treatments directly, which allows us to use statistical techniques to estimate potential clinical utility of the model. In addition, we sought to create a model that relied only on clinical and demographic information to maximize accessibility. We also worked to make our model interpretable at the individual patient level.

## 2 Methods

### 2.1 Data

We analyzed patient-level data from two major trials: CO-MED, by Rush et al. (2011) and the first level of STAR*D by Trivedi et al., 2006. CO-MED enrolled 665 outpatients with non-psychotic depression who were randomized to three treatment arms: escitalopram and placebo, bupropion and escitalopram, or mirtazapine and venlafaxine. The purpose of the trial was to assess whether combination treatment was superior to monotherapy, but similar remission rates were observed in each arm. STAR*D is the largest pragmatic trial of depression treatment to date. In the first of the four levels of the study, all patients were treated with citalopram and the remission rate was 33% (n = 2757, see case selection below). These studies were ideal for our analysis because they included similar outcome measures, the Quick Inventory of Depressive Symptomatology Self-Report (QIDS-SR16); recruited similar patients, those with at least moderately severe depression as determined by the Hamilton Depression Rating Scale (HDRS); included patients treated in both psychiatric and general practice settings; collected similar clinical and demographic information; treated patients for similar lengths of time, 12 weeks in the acute phase of COMED and 12-14 weeks in STAR*D Level 1; and used measurement-based care protocols to adjust doses, i.e. doses were adjusted based on patient symptom scores. Unless otherwise noted, we defined remission as being a score of 5 or less on the QIDS-SR 16.

In STAR*D, our focus was to assess subjects at baseline and predict whether they went into remission after Level 1 treatment was administered (between weeks 2 and 14). We removed 27 subjects who either did not have at least moderately severe depression (i.e., a score of at least 16 on the HRSD questionnaire), did not complete at least one week of treatment, did not return for a visit at week 2, or did not return for a depression assessment in their final week.

We then merged the STAR*D dataset (2,757 subjects) with the CO-MED dataset (665 subjects). We first removed all features that the individual datasets did not have in common. We then merged the datasets, giving us 213 features with a combined 3,222 patients. The combined dataset featured four different treatment groups (three from CO-MED and one from STAR*D) which were labelled by “drug assigned”. This variable ensured that we could calculate differential treatment benefit in our model.

### 2.2 Data Processing and Feature Selection

We employed a variety of methods to ensure the retention of salient features without discarding useful information. Our motivation for this approach is that while redundant features do not add any additional useful information by definition, they do add to the complexity of the modeling and detract from interpretability (Keogh and Mueen, 2017). Our variable curation procedure can be broken down into several steps, each eliminating more and more variables: variance thresholding (removing variables with less than a certain variance), recursive feature elimination with cross validation (RFECV), and feature importance extraction. We were able to identify these optimal parameters for our feature selection by running the features with our neural network and assessing the prediction metrics. Optimistic bias is commonly an issue with thorough feature selection processes. To handle this we extracted our test set of 200 subjects before performing these methods. These methods were implemented from the Python package Sci-kit Learn (Pedgregosa et al., 2011). Figure 1 details this process within the “Phase 1” subheader. Future analyses can use knockoff methods that controls false discoveries (Candes et al., 2018).

**Figure 1.**
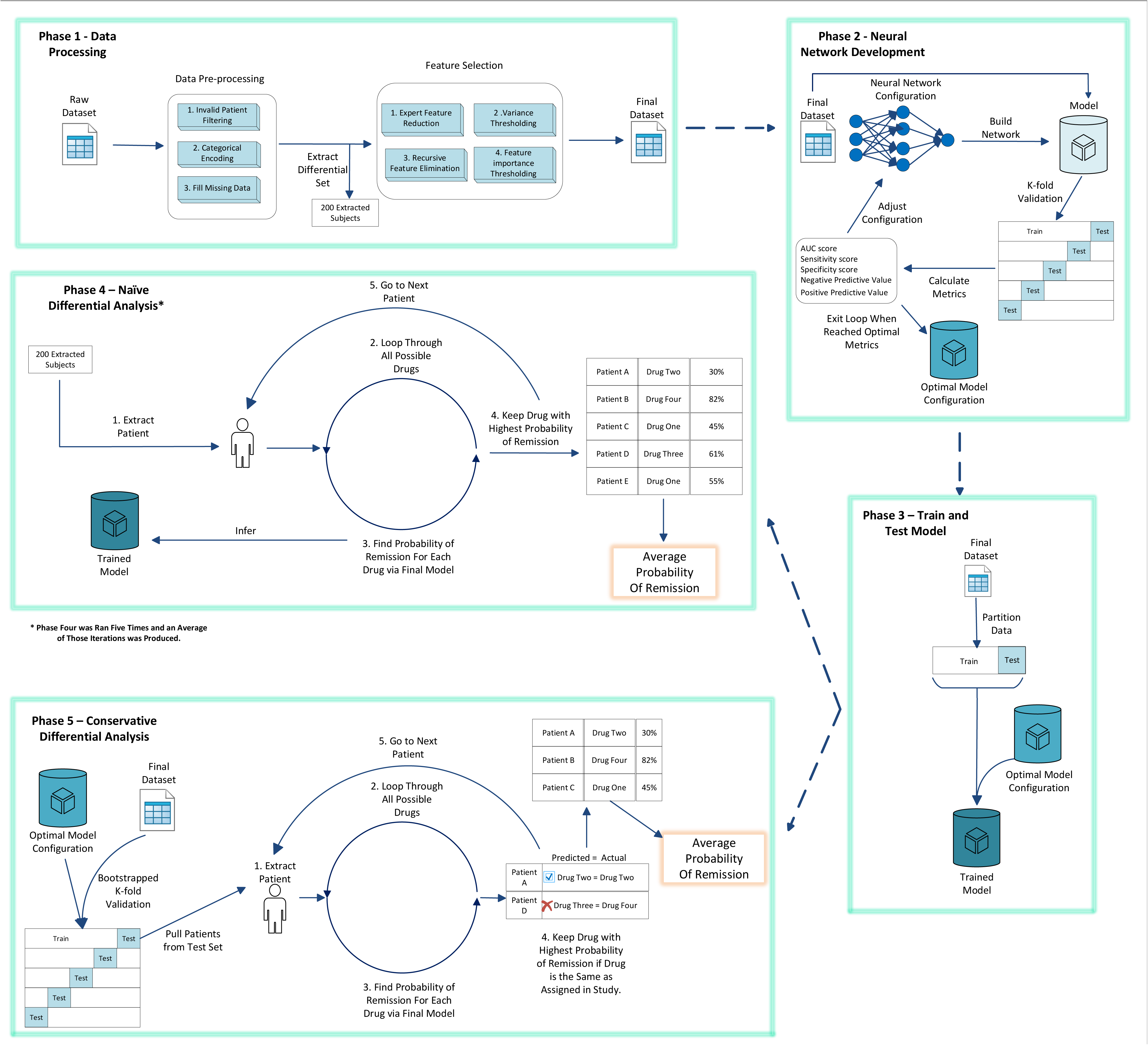
Drug Differential Analysis **Phase 1 (Data Processing)**: The raw data set was preprocessed and then fed through a feature selection procedure to produce a final dataset. Before feature selection occurred, 200 subjects were held out. **Phase 2 (Neural Network Development)**: We configured our neural network and used our final dataset to perform k-fold validation to produce metrics. During training, we iteratively optimized our neural network tuning parameters. **Phase 3 (Model Training and Testing)**: Our dataset was split into three sets: differential analysis set, train set, and test set. The differential analysis set contained 200 patients. To conclude Phase 3 we used our train and test set to train and analyze our model. **Phase 4 (Naïve Differential Analysis)**: We iterated through each subject of our differential analysis set. For each patient we used our neural network with each possible drug to find the probability of remission with that given drug. Once the patient had a probability of remission for each drug, the drug with the highest probability was retained. Once all 200 patients had a probability of remission, we took an average of those probabilities. This process was run five times and the average was computed. **Phase 5 (Conservative Differential Analysis)**: We used k-fold validation with our final dataset. The patients within the test set for each fold were used to perform the differential analysis. Also, to be more conservative we only kept patients if the drug our neural network produced the highest probability of remission for, was the drug they received in the study. We then took the average of all patients kept after all folds from the k-fold validation process.

For the first step, we ran various models under different variance thresholds and found that setting the threshold to 0.2 yielded the best performing model in terms of accuracy of remission prediction. This means that any feature with a variance less than 0.2 across all samples was removed. When testing various thresholds we initially tested and analyzed CO-MED and STAR*D individually to ensure that known relevant features in the literature were not being removed. The 0.2 threshold found was chosen because it was optimal for CO-MED and STAR*D individually, as well as when the datasets were combined. In the second step, the RFECV procedure employs a specified machine learning model to analyze the predictive strength of features independent of one another. We opted for a random forest classifier due to its robustness to hyperparameters (e.g. number of classification trees and the number of variables to try for the split rules in the inner nodes). To further ensure the robustness of the important features selected, and reduce optimistic bias, we used three-fold cross validation. Lastly, we employed feature importance thresholding via randomized Lasso. This procedure randomly shuffled the samples and selected a set of features most closely linked to the label we were trying to predict (i.e. remission). We resampled 200 times and ranked feature importance based on how many times a feature was selected from the previous step. We then removed the features that did not exceed our “importance” threshold of 75%.

Both our CO-MED and STAR*D models had imbalanced classes because the majority of subjects were part of the non-remitter class (only 34% of subjects in STAR*D and 36% of subjects in CO-MED remitted). Due to this class imbalance, we decided to use stratified sampling (Lang et al., 2016) when creating our training and test set for our validation. This ensured that we were training and testing on mutually exclusive but similarly distributed data, preventing bias towards learning and predicting only the majority class. Other techniques like oversampling and undersampling risk information loss (He et al., 2009); therefore, stratified sampling was the best method for our purpose.

### 2.3 Neural Network Architecture and Configuration

Given the structured nature of the data collected in these studies, we opted for fully connected dense neural networks (DNN). To build, train, and evaluate all of our DNN configurations, we employed the open-source package Vulcan (see Software Note) DNNs allow us to capture complex, non-linear relationships likely present in psychiatric data (e.g. mediation and moderation effects, which are unknown a priori). To prevent overfitting, we limited our model’s learning capacity (with the use of a shallow network) in order to explore more of the solution space before finding an optimal location (Caruana, Lawrence, & Giles, 2001). Every node in the network was activated using the scaled exponential linear unit (SELU) function (Klambauer et al., 2017) while the final prediction layer used the softmax function to determine remission probabilities. We one-hot encoded our response variable to create a multi-class problem that uses categorical cross entropy as the optimizer cost function (Goodfellow et al., 2016). We used the Adam optimizer for learning the network parameters (Kingma & Ba, 2014) with a learning rate of 0.0001. To further help with model generalization, we used a 50% dropout rate (Srivastava et al., 2014). During the evaluation phase, we used 10-fold cross validation. Each fold allowed the model to train for 200 epochs.

## 3. Results

### 3.1 Differential Treatment Efficacy Prediction

In our differential treatment prediction analysis, we conducted two separate experiments to determine the relative increase in population remission rates when using the treatment recommended by our model compared to the treatment patients were assigned in the study. These experiments were done on the combined CO-MED and STAR*D datasets using the same trained DNN model. This model was built using our feature curation procedure (sec. 2.2) and consisted of 17 features, listed in Table 1. After DNN training, this model yielded a 0.69 macro^1^ AUC, 0.70 macro PPV, 0.70 macro NPV, 0.72 macro sensitivity, 0.72 macro specificity. We used our 200 held-out subjects to perform external validation before running our differential analysis.

**Table 1.**

| The 17 Most important features in the Differential treatment prediction model |
| --- |
| Number of years in formal education |
| Monthly household income |
| HAM-D Somatic energy |
| Bothered by aches/pains |
| QIDS Weight (increase) last 2 weeks |
| QIDS Mood (sad) |
| QIDS Suicidal ideation |
| QIDS Total Score |
| QIDS Energy/Fatigability |
| QIDS Sleep Onset Insomnia |
| Jumpy because of a Trauma |
| Ever Witnessed a Traumatic Event |
| Did reminders of a traumatic event make you shake, break out into a sweat, or have a racing heart. |
| How many hours actually worked |
| Anxiety being in crowded places |
| Eat a lot when not hungry |
| Drug Assigned |

Our first “naïve” experiment was aimed at estimating the performance of the model if it was applied blindly to a “new” clinical population. To create a test set, we isolated 200 subjects from the dataset, ensuring that these were split to reflect the distribution found in the original datasets. This meant that 20% of subjects were from the CO-MED study and 80% were from the STAR*D study. We then trained our neural network on the remaining subjects. In order to get a probability of remission, we passed each subject in the held-out sample four times through the final model, once for each possible drug in the dataset. The output of the forward pass was the probability of remission for that subject for that given drug. For each of the 200 subjects we took the drug with the highest probability of remission and obtained the mean remission rate of the 200 subjects. Finally, we took the difference between the mean remission rate of the entire dataset and our mean remission rate for the 200 test subjects. We ran this test five times and took the average of those five tests which produced an improvement of ~11% in the population remission rate from 34.33% to 46.12%, a relative improvement of over 30%, when using our neural network to assign patients to drugs when compared with the baseline study drug assignation.

In this naïve version of the analysis, we looked at hypothetical cases in which we did not necessarily know the outcome of giving a certain patient a drug. Our second “conservative” analysis only considered the non-hypothetical cases in which we knew the outcome of giving a specific treatment option to a patient. This second analysis was inferential, and details of its methods is found in Kapelner et al. (2017). This analysis answered the question: does our DNN personalization model beat a null model --- does it improve patient outcomes more than chance allocation? For this null allocation, we used 1,000 independent and identically distributed bootstrap samples. In creating these new samples, we resampled from the original dataset with replacement, trained a model several times through 10-fold cross validation, and computed an “improvement score” that compares the chance remission rate with the improved remission rate found using the personalized assignations from our DNN model. To exclude hypothetical cases, we compared improvement scores between patients who actually received (by chance) the drug our model predicted they should have received to the rest of the study population, who were allocated to a drug as per study protocol. This inferential approach is extremely conservative in a retrospective analysis for two reasons. First, we only consider patients who actually received the treatment recommended by the predictive model. Second, our sample size is greatly reduced as we only predict on data that we did not use to build the DNN, i.e. only 10% of the data. We observed a significant improvement of 2.5% (p-value of 0.01, C.I. 2.48% ± .5%) from 34.33% to 36.8% population remission rate, a relative improvement of 7.2%.

## 4 Discussion

We used deep learning to create a personalized medicine model that is useful as a proof-of-concept for differential treatment selection. We combined data from the CO-MED and STAR*D datasets to produce a pooled dataset with four different treatment types. Our main goal was to demonstrate that our model was capable of performing differential treatment selection, and to estimate improvement in patient outcomes. We produced a well-validated model via cross-validation within the test set, validation on a held-out dataset, and only then applied the model to our held-out sample of 200 sub-sampled but mutually exclusive patients. By retaining the treatment selection (the drug class) as a feature in our model, we could then generate predictions for each drug class for all 200 held-out patients. With these predictions, we estimated the remission rate for these patients had they been assigned a drug based on our model, showing an 11% average increase in population remission rates. This potential remission increase of 11% is clinically significant, essentially increasing remission rates by a third over measurement-based care alone in STAR*D and CO-MED. We then further validated our results by proving statistically that our personalized models beat chance remission via a conservative bootstrapping procedure, which demonstrated a significant increase in overall remission rate of 2.5% (p = 0.01). The conservative analysis’ bootstrap estimate of out-of-sample improvement is only valid asymptotically and also has a large standard error. Thus, the population remission improvement estimated using this analysis does not accurately depict the likely effect size of the differential prediction benefit. Rather, it represents more of a “lower bound” of sorts. The true benefit will likely lie between the numbers produced by the conservative and naive analyses.

A critical contribution of our work is extending these evaluation metrics to models that cover a number of treatment options. This is significant because most previous literature has focused on helping to select between two treatment options, when in clinical practice clinicians are faced with over a dozen treatment choices. Models that can help select between a number of different treatments may have significant clinical impact when properly validated and implemented.

As can be seen in the supplementary material, a version of our model trained only on STAR*D generalizes to all three arms of CO-MED, in contrast to the model by Chekroud et al., which only generalized to two of the three arms. Importantly, there was significant overlap between the 25 features included in the Chekroud et al. model and the 14 included in our own (Supplementary Table 1). This initial analysis demonstrates that deep learning can provide improved results when compared to other machine learning techniques while potentially using more parsimonious feature sets allowing for greater ease of implementation in clinical settings. Deep learning has often been found to significantly outperform other machine learning techniques as the size of the dataset increases (L’Heureux et al., 2017), and as such finding even a small advantage for deep learning in this fairly small (by deep learning standards) dataset leads us to speculate that deep learning will perform significantly better than other techniques on larger datasets.

We examined the population remission rate, and as such it is difficult to determine the benefit for each individual patient. Future analyses, along the lines of the Personalized Advantage Index (PAI) (Derubeis et al., 2014) may be able to help estimate the individual benefit with model-predicted treatment. However, what is intriguing is that despite equal effectiveness of these treatments at a population level, we are able to use individual patient differences in predicted remission generated by varying the assigned drug to improve the overall remission rate. If all treatments were truly equally effective for all individuals, we could have expected a model that approximated but did not improve upon the remission rate. The finding of a projected improvement supports a personalized medicine approach based on individualized prediction of response to treatment.

Deep learning has often been labelled as a “black box”, meaning deep learning models can be a challenge to interpret (Samek et al., 2017). Interestingly, when using only clinical and demographic measures we find that deep learning systems provide a list of features which could be interpreted by clinicians. As demonstrated in Box 1, the most important features used in the prediction for each individual patient can be recovered, providing insights personalized to that patient. This ‘personalized prediction report’ demonstrates that deep learning-based tools may be able to provide information that is useful for understanding individual clinical cases.

It is interesting to note that there are two categories of features in Table 1: those likely to predict overall probability of remission and treatment-specific features. For example, level of education and income - which have both been found in other studies of remission prediction (Carter et al., 2012) - are both unlikely to be specifically related to any one drug’s mechanism-as opposed to sleep pattern, which may be relevant to a particular drug’s mechanism. It is important to review the features that have been identified in our model and compare them to models in previously published work, and a full discussion is included in the Supplementary Materials. Specific symptoms retained by our model may contribute to differential prediction. For example, in the Individualized Patient Reports (Box 1) Patient B was predicted to do better with a combination of Bupropion and Escitalopram and one of the five most important features in that prediction was a tendency to eat a lot when not hungry, which is interesting given the fact that Bupropion is often used clinically in cases of hyperphagia or when weight gain is to be avoided (Patel et al., 2016; Anderson et al., 2002).

Beyond the demonstration of a differential treatment prediction tool for n > 2 treatments, we also provide evidence, in accordance with recent papers (Lin et al., 2018), that deep learning can be readily applied to psychiatric datasets. This is an important development in the field as deep learning is well suited to the analysis of multimodal data and may help bring together data from neuroimaging, genetics, or wearable technology with clinical and demographic measures; this is another reason for developing sound methodology for working with deep learning in psychiatry.

It remains to be seen if the effect we estimate materializes in a clinical environment. That being said, our model’s AUC of 0.69 is potentially clinically significant because of the low baseline response rate to antidepressants. Any tool that could help improve the accuracy with which clinicians can identify which patients are likely to benefit from which treatments may be welcomed by the clinical community. This is especially true given the low risk of choosing ‘wrong’ (i.e. choosing an ineffective first line antidepressant, which occurs commonly) and the high reward of choosing ‘right’-choosing the right medication for the individual patient and reducing time to remission. In this context a model with a similar performance to our own might be clinically significant because it likely will outperform the status quo without significantly increasing the risk of adverse events. In addition, it is worth considering the model in terms of potential benefit to the population. Using our model, we can estimate that between 80 (based on the conservative analysis) and 354 (based on the naive analysis) more patients in the dataset would have reached remission after a single trial of pharmacotherapy.

We note limitations to our study. In our pooled dataset, patients assigned citalopram vastly outnumbered those prescribed other treatments, limiting the extent to which we can be confident that we were predicting differences between four completely distinct treatment classes. Further studies analyzing datasets with smaller class imbalances are necessary. We recognize that while CO-MED was a randomized trial, all patients in STAR*D were initially assigned to citalopram. However, given the very similar patient populations and study protocols, we felt justified in combining the studies as if they were arms of the same study.

## 5 Conclusion

We have demonstrated proof-of-concept of using a deep learning model to predict remission and to guide differential treatment selection for n > 2 treatments. More data is required to validate these methods for other treatment types to create a clinically useful tool. Furthermore, clinical trials are needed to determine if our hypothetical treatment assignment is translatable to real patients. The flexibility and versatility of our model supports the idea that deep learning will be a useful technique going forward in the field of personalized medicine.

### Box 1

#### Examples of differential predictions for two subjects (“Personalized Prediction Reports”)

Here we show ‘reports’ for two patients showing the drug they received in the study and whether or not they went into remission; the drug the network predicted and the predicted remission rates for that drug and the drug the patient actually received; and the five most important features for that patient based on the neural network (all features were used for both patients, but the five listed were weighted more by the network for that patient)

Subject A was originally given Escitalopram and went into remission. Our neural network found Escitalopram to have an 87.57% chance of remission. The five most important features for this patient and the attendant responses were:

- Total monthly income? 2,200
- How many years of formal education? 12
- Experienced weight increase within the last two weeks? Has gained 5 pounds or more,
- Have you been feeling down, blue, sad or depressed? Feels sad less than half the time
- Do you eat a lot when not hungry? No

Subject B was originally given Citalopram and did not go into remission. Our neural network found Citalopram to have a 38.03% chance of remission, while Bupropion SR & Escitalopram had a 60.43% chance of remission, indicating that changing the medication may have been beneficial for this patient. The five most important features for this patient and attendant responses were:

- Have you ever witnessed a traumatic event? No
- Did reminders of a traumatic event make you shake, break out into a sweat, or have a racing heart? No
- Have you had any trouble falling asleep when you go to bed? Takes at least 30 minutes to fall asleep, more than half the time.
- Have you been feeling down, blue, sad or depressed? Feels sad less than half the time
- Do you eat a lot when not hungry? Yes

#### Software Note

The Vulcan platform used for this work is open source and can be found here: (https://github.com/Aifred-Health/Vulcan).

## Supporting information

Supplementary word document

## Author Contribution

JM, RF, and DB contributed equally to this paper and are considered co-first authors. JM programmed and ran the analyses and contributed to the Vulcan pipeline and created Figure 1 and the Tables. RF built the Vulcan pipeline and oversaw the analyses and helped conceptualize the analyses and edited Figure 1. DB conceptualized the analyses, wrote and edited the manuscript. AK conceptualized the conservative analysis and assisted in the conceptualization of the other analyses, and edited the manuscript. KP, ES, and SI assisted with background research and editing the manuscript. MM provided input into the analyses and evaluation metrics. GT supervised the work, helped secure access to the data, and edited the manuscript.

## Ethics Approval

This work was approved by the Research Ethics Committee of the Douglas Mental Health University Institute

## Role of the Funding Source

No funding was provided to AK and GT. No funding was secured for this study in particular, but JM, RF, DB, KP, ES, MM and SI are all employees, contractors, or shareholders in Aifred Health and were compensated monetarily or with stock by Aifred Health while they were working on this paper. This compensation was not tied to the results of this paper. This paper directly supports the work of Aifred Health, which is developing decision support tools for treatment selection in depression.

## Footnotes

1 Macro metrics are calculated by calculating the metric for each class and then averaging these

