## Supplementary word document for "Differential Treatment Benefit Prediction For Treatment Selection in Depression: A Deep Learning Analysis of STAR*D and CO-MED Data"

**Replication of Previous Results and Advantage of Deep Learning**

We began with a replication of the Chekroud et al. 2016 results- that is, we trained a model on the citalopram data from STAR*D only and then tested on the independent data from CO-MED. The results can be found in Supplementary Table 2 and demonstrate that the deep learning model achieved an AUC of 0.7 on the citalopram dataset and was able to generalize to all three arms of CO-MED, using a model with only 14 input features. Critically, as will be seen in the discussion, the model recovered some of the same input features as the Chekroud model. The fact that we were able to produce a deep learning model with a generalizable solution and which was able to replicate previous data was an important check of the applicability of deep learning models to psychiatric datasets of this size and composition. We then proceeded to produce a model using the combined dataset described above; this model is the focus of the main paper.

With respect to data collection, the deep learning model’s smaller feature set and lower reliance on clinician-collected data from the HAM-D may help reduce the burden of data collection in clinic. For example, while both models relied on the total score from the 16 item QIDS, our deep learning replication of the Chekroud model only required 10 non-QIDS questions, for a total of 26 questions, only one of which needed to be asked by a clinician. In contrast, the Chekroud model would have required a total of 47 questions, including 17 that needed to be asked by a clinician (all questions in the HAM-D). Given that the time a doctor has to spend with a patient is often limited, reducing the data-collection burden on clinicians is potentially significant when it comes to implementing treatment selection algorithms in the clinic.

**Supplementary Table 1.** The most salient features for remission prediction from our analysis compared against features from Checkroud et al. 2016.

| Most important features (STAR*D, generalizing to COMED) (14) | Chekroud et al. 2016 most important features (25) |
| --- | --- |
| Number of years in formal education | Years of education |
| Anxiety being home alone | QIDS Psychomotor agitation |
| Anxiety being in crowded places | QIDS Energy or fatigability |
| Anxiety standing in long lines | Black or African American |
| HAM-D Somatic energy | HAM-D Somatic energy |
| Bothered by aches/pains | Currently employed |
| QIDS total severity | Initial QIDS total severity |
| Jumpy because of a trauma | HAM-D somatic anxiety |
| Monthly household income | HAM-D loss of insight |
| Ever witnessed traumatic event | Initial HAM-D depressive severity |
| QIDS Mood (sad) | QIDS mood (sad) |
| QIDS Suicidal ideation | Did reminders of a traumatic event make you shake, break out into a sweat, or have a racing heart. |
| QIDS Weight (increase) last 2 weeks | HAM-D Delayed insomnia |
| How many hours actually worked | Have you ever witnessed a traumatic event such as rape, assault, someone dying in an accident, or any other extremely upsetting event? |
|  | Did you try to avoid activities, places, or people that reminded you of a traumatic event? |
|  | White |
|  | Did any of the following make you feel fearful, anxious, or nervous because you were afraid you’d have an anxiety attack in the situation? Standing in long lines. |
|  | Did any of the following make you feel fearful, anxious, or nervous because you were afraid you’d have an anxiety attack in the situation? Driving or riding in a car |
|  | Have you been bothered by aches and pains in many different parts of your body. |
|  | HAM-D suicide |
|  | Depressed mood most of the day, nearly every day. |
|  | Did you have attacks of anxiety that caused you to avoid certain situations or to change your behaviour or normal routine? |
|  | Ever taken sertraline |
|  | Number of previous major depressive episodes |
|  | QIDS sleep onset insomnia |

##### **Supplementary Table 2**. Evaluating internal and external generalisability of our STAR*D trained model which was then tested on each arm of CO-MED, compared to Chekroud et al 2016. Note: The Checkroud Model was evaluated on the metric of accuracy in the generalizability analyses.

| Dataset tested | AUC | NPV | PPV | Sensitivity | Specificity |
| --- | --- | --- | --- | --- | --- |
| STAR*D (14 input features) | .70 | .64 | .64 | .60 | .60 |
| STAR*D - Chekroud et al (25 input features) | .70 | .653 | .640 | .628 | .662 |
| CO-MED (Escitalopram & Placebo arm) -  (14 input features) | .64 | .58 | .58 | .59 | .59 |
| CO-MED (Escitalopram & Placebo arm) - Chekroud et al (25 input features) | .596 (accuracy) | .560 | .65 | .494 | .708 |
| CO-MED (Bupropion SR & Escitalopram arm) (14 input features) | .63 | .60 | .60 | .61 | .61 |
| CO-MED (Bupropion SR & Escitalopram arm) - Chekroud et al (25 input features) | .597 (accuracy) | .597 | .597 | .561 | .632 |
| CO-MED (Venlafaxine & Mirtazapine arm) (14 input features) | .63 | .60 | .60 | .61 | .61 |
| CO-MED (Venlafaxine & Mirtazapine arm) - Chekroud et al (25 input features) | .514 (accuracy) | .50 | .539 | .389 | .647 |

##### **Extended Discussion of Retained Features**

A solid understanding of the features in play will be key in the elaboration of a personalized medicine approach. In our 17 feature model, we found many features that were similar to those identified by Checkroud et al., 2016. In terms of demographics, both models identified the number of years in formal education as being predictive of response. In terms of symptoms, total symptom scores, suicidality, sleep disturbance, somatic symptoms, and energy and fatigability were common between both models. It is interesting to note that Iniesta et al., 2018, also found that initial severity, sleep changes, somatic symptoms, and fatigability were predictive of escitalopram response. This coheres well with literature that notes the importance of somatic symptoms and fatigue, which are often residual symptoms, in achieving and maintaining remission (Papakostas et al., 2004; Fava et al., 2014) , as well as with literature noting that those with a higher initial severity of depression are less likely to remit (De Carlo, Calati & Serretti, 2016; Papakostas & Fava, 2008). Trauma, and current symptoms related to past trauma, were also important features in both our model and the Chekroud model, though these did not appear in the Iniesta model. Trauma has long been recognized as playing a role, likely via genetic, epigenetic as well as psychological mechanisms (Penza, Heim & Nemeroff, 2003; Heim et al., 1997; Klengel et al., 2014; Mann & Currier, 2006), in the pathophysiology of major depression; the fact that trauma does not appear in the Iniesta analysis may be, speculatively, because it is captured in the genetic features which were key to that analysis or because of differences in sample characteristics. Race did not feature in our model, though it was present in the Chekroud model. This is interesting because our model also retained level of income, which was not present in the Chekroud model, given that race and income are often correlated because of discrimination and location (Shapiro, Meschede & Osoro, 2013; Campbell & Kaufman, 2006); some of our future work will focus on elucidating this finding. Anxiety, specifically of an agoraphobic type (anxiety being in crowded places), was also found in both the Chekroud model and our own, which is consistent with literature noting that patients with anxiety are less likely to respond to antidepressants (Iosifescu, Bankier & Fava, 2004; Papakostas & Fava, 2008, Turecki & Berlim, 2007). Finally, it is interesting to note that the Iniesta model found that interest-activity (Uher et al., 2012) was a predictive factor for determining escitalopram response; both our model and the Chekroud model noted energy and fatigability and employment as important factors, with our model specifically noting the actual number of hours of work people engaged in (and not just their employment status, which can be less reliable an indication of activity if a patient is off-work) as being important.

Models incorporating genetic information are gaining in importance, such as that produced by Iniesta et al., in 2018 which was able to predict remission in patients taking escitalopram with a model using clinical and genetic features that achieved an AUC of 0.77, and in patients taking notryptiline using a model with only genetic factors with that achieved an AUC of 0.77; further work using deep learning in this area would do well to include genetics as a multi-modal input. An interesting model design would be one that could use genetic information if it is available, which would make the model adaptable to different resource contexts.

##### **Supplementary Table 3**. Additional model evaluations with STAR*D and CO-MED

This table demonstrates our network trained and tested on CO-MED alone, along with the full results of our 17 feature combined STAR*D and CO-MED model.

| Dataset that the model was trained on | AUC | NPV | PPV | Sensitivity | Specificity |
| --- | --- | --- | --- | --- | --- |
| CO-MED internal validation w/ 25 input features | .80 | .704 | .704 | .72 | .72 |
| STAR*D + CO-MED shuffled internal validation (17 input features) | .69 | .64 | .64 | .60 | .60 |

##### **Supplementary Table 4. Evaluating combined STAR*D/CO-MED prediction with baseline Random Forest and Logistic Regression models.**

| Model | AUC | NPV | PPV | Sensitivity | Specificity |
| --- | --- | --- | --- | --- | --- |
| Random Forest Classifier  (17 input features) | 0.67 | 0.63 | 0.63 | 0.58 | 0.58 |
| Logistic Regression Classifier (17 input features) | 0.68 | 0.65 | 0.65 | 0.61 | 0.61 |

There is significant debate in the literature about appropriate trade-offs between algorithmic power and interpretability. Deep learning has often been labelled as a “black box”, meaning deep learning models can be a challenge to interpret (Samek et al., 2017), leading many to assert that a significant improvement in predictive power over other techniques would be necessary to justify its use in the clinical setting. Indeed already in this dataset our deep learning approach outperforms a random forest approach which we found achieved an AUC of 0.67, and performs as well, when predicting the STAR*D Level 1 results, as the boosted regression approach in Checkroud et al., 2016 while using less information and with improved generalization. Deep learning has often been found to significantly outperform other machine learning techniques as the size of the dataset increases (L'Heureux et al., 2017), and as such finding even a small advantage for deep learning in this fairly small (by deep learning standards) dataset leads us to speculate that deep learning will perform significantly better than other techniques on larger datasets.

**Supplementary Table 5: Mean Individual Differences in Remission Likelihood in the Naive Analysis**

|  | Mean Individual Difference | Standard Deviation |
| --- | --- | --- |
| Test 1 | 12.21 | 20.9 |
| Test 2 | 8.78 | 18.32 |
| Test 3 | 9.15 | 17.13 |
| Test 4 | 8.29 | 18.14 |
| Test 5 | 11.39 | 19.87 |

This table shows the mean individual differences in remission likelihood in five re-drawn samples in the Naive Analysis with our model. This allows for a sense of the impact of the algorithm and its variability with respect to projected remission rate improvement.

**Predicting CO-MED remission likelihoods**

We also ran models predicting remission for CO-MED subjects alone (i.e. within the COMED study). We focused on the acute phase (12 weeks). There were 731 subjects screened for the study, but we only used the 665 subjects who entered the study during week 0 and were assigned treatment for our analysis. We calculated remission based on the QIDS-SR features provided. A patient was considered in remission if they had a QIDS-SR score of less than 6 and 8 on their two most recent assessments, respectively (as defined in the study). Our baseline model uses 25 features after feature selection when predicting remission. When predicting remission for our CO-MED model using our feature engineering techniques and stratified samples described above we obtained 0.80 macro AUV, 0.704 macro PPV, .704 macro NPV, 0.72 macro sensitivity, and 0.72 macro specificity. This is demonstrated in the above table.

**Biomarkers:**

Biomarkers that could be incorporated into deep learning modes include electroencephalography (EEG), magnetic resonance imaging (MRI) (Hunter et al., 2007; Wise et al., 2014), biofluids (Gadad et al., 2018) and personal technological devices (Torous et al., 2017). Furthermore, Bradley et al. (2018) showed that using a genetic screening test helped improve remission rates compared to a control group that did not have access to the tests.
